# Characterization, structure and inhibition of the human succinyl-CoA:glutarate-CoA transferase, a genetic modifier of glutaric aciduria type 1

**DOI:** 10.1101/2024.02.07.578422

**Authors:** Susmita Khamrui, Tetyana Dodatko, Ruoxi Wu, João Leandro, Amanda Sabovic, Sara Violante, Justin R. Cross, Eric Marsan, Kunal Kumar, Robert J. DeVita, Michael B. Lazarus, Sander M. Houten

**Author notes:** co-corresponding authors, Correspondence Michael B. Lazarus, Department of Pharmacological Sciences, Icahn School of Medicine at Mount Sinai, New York, New York 10029, United States.; Sander M. Houten, Department of Genetics and Genomic Sciences, Icahn School of Medicine at Mount Sinai, New York, New York 10029, United States. Equal contribution (Susmita Khamrui, Tetyana Dodatko, Ruoxi Wu, João Leandro, Amanda Sabovic). Drug Discovery Institute, Icahn School of Medicine at Mount Sinai, New York, NY 10029, USA.

## Abstract

Glutaric Aciduria Type 1 (GA1) is a serious inborn error of metabolism with no pharmacological treatments. A novel strategy to treat this disease is to divert the toxic biochemical intermediates to less toxic or non-toxic metabolites. Here, we report a novel target, SUGCT, which we hypothesize suppresses the GA1 metabolic phenotype through decreasing glutaryl-CoA. We report the structure of SUGCT, the first eukaryotic structure of a type III CoA transferase, develop a high-throughput enzyme assay and a cell-based assay, and identify valsartan and losartan carboxylic acid as inhibitors of the enzyme validating the screening approach. These results may form the basis for future development of new pharmacological intervention to treat GA1.

## Introduction

Succinyl-CoA:glutarate-CoA transferase (SUGCT) is a mitochondrial dicarboxyl-CoA:dicarboxylic acid CoA transferase originally purified from rat liver mitochondria [1–3]. *C7orf10*, the gene causal for glutaric aciduria type 3 (GA3), was later demonstrated to encode SUGCT [4]. Although multiple dicarboxylic acids and their CoA esters can serve as substrates, this genetic association suggests that the succinyl-CoA-dependent conversion of glutaric acid into succinate and glutaryl-CoA is the main physiological reaction (**Fig. S1A**). SUGCT belongs to the type-III CoA transferase (CaiB/BaiF) superfamiliy of proteins, which includes CoA transferases as well as racemases. Structures have been solved for some family members [5–11], but no eukaryotic structures including human SUGCT have been reported previously.

The main biochemical abnormality of GA3 patients and SUGCT deficient mice is increased urine levels of glutaric acid [4, 12, 13]. Glutaryl-CoA is an intermediate of lysine degradation. It is thought that a significant fraction of glutaryl-CoA undergoes spontaneous or enzyme-mediated hydrolysis to glutaric acid [14, 15], and SUGCT function is needed for re-esterification to glutaryl-CoA before further metabolism by glutaryl-CoA dehydrogenase (GCDH) can occur. Since glutaric acid would otherwise be a dead-end metabolite excreted in urine, this process is viewed as an example of metabolite repair [16]. Expression of SUGCT is not uniform, with high mRNA levels in kidney and liver.

GA3 is considered a biochemical phenotype with limited or no clinical significance [4, 17–19]. In practice, this condition can be diagnosed through biochemical and genetic methods, but is thought to be not harmful. GA3 has been reported in up to 18 individuals; many are asymptomatic, whereas others have symptoms such as gastrointestinal problems and cyclic vomiting [4, 17, 19–24]. Encephalopathic crises, which are a characteristic feature of glutaric aciduria type 1 (GA1), have not been reported in GA3 patients. Based on several observations, there is an uncertain causal relationship between clinical disease and the biochemical diagnosis of GA3. First, healthy cases have been identified in urine-based newborn screening as well as studies of families investigated for a symptomatic case of GA3 or an unrelated diagnosis [4, 17, 19, 23]. Second, the clinical symptoms associated with GA3 are non-specific, further increasing the possibility that the biochemical abnormalities were associated by chance. This implies that the clinical symptoms in these patients are caused by other genetic or environmental factors. Indeed, in some cases alternative genetic causes were identified that may better explain the clinical disease, including β-thalassemia, 6q terminal deletion syndrome, autoimmune hyperthyroidism and trichothiodystrophy type 4 [17, 19–21]. Some of the GA3 cases were followed for more than 15 years and remained asymptomatic [4, 17, 23]. Last, the p.Arg336Trp

variant that causes GA3 in the Amish (rs137852860 [4]) has an allele frequency of 9.3% in the Amish and 0.8% in non-Finnish Europeans and the gnomAD database contains 10 individuals homozygous for this variant. The presence of homozygotes in this population-based database is generally considered good evidence that a variant is not disease-causing [25]. Therefore, the current consensus in the field is that GA3 due to SUGCT deficiency does not lead to a clinically significant disease [18]. The association of GA3 with clinical symptoms is likely due to ascertainment bias. Similarly, SUGCT deficiency in mice does lead to a recognizable clinical phenotype [12, 13].

In contrast to GA3, GA1 (MIM 231670) is a serious neurometabolic disorder. If left untreated, GA1 patients will likely develop a complex movement disorder due to striatal injury after an acute encephalopathic crisis, or with insidious onset [26, 27]. These crises typically occur in the first year of life and are often triggered by a catabolic state such as those that occur during childhood illnesses. GA1 is caused by mutations in *GCDH* [28–30] leading to a deficiency of GCDH and the accumulation of neurotoxic levels of metabolites including glutaric acid and 3-hydroxyglutaric acid. GA1 has been included in newborn screening programs in the US and around the world because patients benefit from immediate intervention [31]. Current treatment is suboptimal and therefore GA1 represents an unmet “orphan” medical need. We have previously shown that in a mouse model, naturally occurring variation in *Sugct* is a modifier of the biochemical phenotype of GA1 [12]. SUGCT deficiency in GA1 mice decreases the accumulation of 3-hydroxyglutaric acid and glutarylcarnitine (C5DC, **Fig. S1B**) [12]. Therefore, we have proposed that inhibition of SUGCT will decrease the neurotoxic metabolite buildup in GA1 and serve as a novel therapeutic target for this disease. In order to facilitate the development of SUGCT as a pharmacological target, we report novel in vitro and cell assay for SUGCT activity, the structure of SUGCT and the identification of the first SUGCT small molecule inhibitors.

## Materials and Methods

### Materials

A full length wild-type *SUGCT* cDNA cloned into pCR-BluntII-TOPO (BC098117, identical to NM_001193313.2) was obtained from Transomic (Huntsville, AL). The Flp-In Core System included the pOG44 and pcDNA5/FRT plasmids (Thermo Fisher Scientific). These plasmids enable stable expression of a cDNA under control of the CMV promoter through integration into a host genome via Flp recombinase-mediated DNA recombination at an FRT site. A Flp-In 293 cell line containing a single stably integrated FRT site at a transcriptionally active genomic locus was purchased from Invitrogen (R75007, Thermo Fisher Scientific). The FDA approved drug library was from APExBIO (L1021, Houston, TX). For immunoblotting, we used a rabbit polyclonal SUGCT antibody, which is unfortunately discontinued (NBP2-69820, Novus Biologicals) [12]. Valsartan, losartan carboxylic acid and other Angiotensin II Receptor Blockers were obtained from Cayman Chemicals. Valsartan ethyl ester was obtained from MedChemExpress. 3-hydroxy-3-methylglutaric acid (HMG) was obtained from TCI or Cayman Chemicals.

### Cloning procedures and generation of stable SUGCT overexpressing 293 cells

The *SUGCT* coding sequence was amplified by PCR by using PrimeStar GXL polymerase (Takara Bio Inc) using the following primers (start and stop codon in bold): forward 5’-GATATA <u>GGTACC</u> **ATG** CTG GCG ACG CTG-3’; reverse 5’-GTGGCC <u>CTCGAG</u> **TCA** GTG AGT TTC ATG TTG G-3’ and then purified using the Monarch PCR plus DNA purification kit (NEB). The purified PCR product was digested by KpnI and XhoI (NEB, underlined in primer sequences) and repurified. The pcDNA5/FRT vector was also digested by KpnI and XhoI followed by purification. The *SUGCT* insert and pcDNA5/FRT vector were ligated by using the Quick Ligation Kit (NEB) and transformed to TOP10 chemically competent cells (Thermo Fisher Scientific). Successful cloning was confirmed by sequencing. pcDNA5/FRT-SUGCT and pOG44 plasmid DNA was purified by NucleoBond Xtra Midi Plus EF DNA purification Kit (Macherey-Nagel). Purified pOG44 and pcDNA5/FRT-SUGCT plasmids were co-transfected into Flp-In 293 cells at a 9:1 ratio by using Lipofectamine 2000 reagent (Invitrogen, Thermo Fisher Scientific). Flp-In 293 cells were cultured in DMEM supplemented with 10% FBS, Pen/Strep and 100 µg/ml Zeocin. For transfection, the culture medium was removed and cells were washed with warm PBS, followed by addition of DMEM with 10% FBS and Pen/Strep but without Zeocin. Plasmid DNA Lipofectamin 2000 complexes were made in Opti-MEM according to manufacturer’s protocol and added to this medium. Transfected cells were incubated at 37°C with 5% CO_2_ for 24h after which the transfection medium was replaced by fresh DMEM supplemented with 10% FBS and Pen/Strep and cells were incubated for another 24h. Cells were then split to 25% confluence and Hygromycin B selective DMEM was added (10% FBS, Pen/Strep and 100 µg/ml of Hygromycin B). Cells were incubated for 2-3 weeks at 37°C with 5% CO_2_. The selective DMEM was replaced every 3-4 days. After Hygromycin B resistant foci were formed, cells were transferred to new flasks and grown in Hygromycin B selective medium until they were 75-80% confluent. Cell pellets were collected for the confirmation of SUGCT overexpression.

### Generation Flp-In 293 cell lines with stable SUGCT overexpression and GCDH KO

*GCDH* was targeted using CRISPR-Cas9 genome editing. For this, a pCas9-GFP plasmid with a previously validated *GCDH* guide sequence was transfected into Flp-In 293 (clone 25 for stable SUGCT overexpression) using Lipofectamine 2000 [32]. After 48h, transfected cells were collected and GFP positive single cells were sorted into a well of a 96 well plate with 50% conditioned DMEM (DMEM collected from the same cell line when cells were 60-90% confluent mixed 1:1 with fresh DMEM supplemented with Pen/Strep and 20% FBS). After 12-15 days of incubation at 37°C and 5% CO_2_ clones appeared in several wells. Clones were first transferred to 24-well plates and after reaching 70-90% confluence were transferred to T25 flasks. Cell pellets were then collected for confirmation of *GCDH* KO through Western blot and DNA sequencing. Overexpression of SUGCT in *GCDH* KO clones was reconfirmed by Western blot.

### LC-MS analysis

The intracellular metabolites were immediately extracted by adding 1 mL ice-cold 5:3:2 extraction solvent (methanol:acetonitrile:water) directly to the plate with cells and incubated overnight at -80°C to aid protein precipitation. The following day, the cells were scraped, and the extracts centrifuged at 20,000xg for 20 min at 4°C. The supernatant was transferred to a new tube and evaporated to dryness in a vacuum concentrator (GeneVac). The samples were reconstituted in 50 µL of 50:50 water:methanol and incubated on ice for 20 min, vortexing every 5 min to ensure adequate re-suspension. All samples underwent one final centrifugation step (20,000xg for 20 min at 4°C) to remove any residual particulate. The reconstituted samples were subjected to LC-MS acquisition using an Agilent 1290 Infinity LC system equipped with a quaternary pump, multisampler, and thermostatted column compartment coupled to a 6546 Q-TOF system (Agilent Technologies) using a dual Agilent Jet Stream source for sample introduction. Reverse phase LC–MS analysis was performed in positive ionization mode using an Acquity UPLC HSS-T3 C18 1.8 um, 100 x 2.1 mm µm particle size (Waters), and using a gradient of solvent A (25 mM formate + formic acid, pH 2.9) and solvent B (100% acetonitrile). The analytical gradient was 0-1 min, 50% B; 1-12 min, 99.5% B; 12-13.9 min, 99.5% B; 13.9-14 min, 50% B. Other LC parameters were as follows: flow rate 400 µL/min, column temperature 40°C, and injection volume 5 µL. MS parameters were as follows: gas temp: 150°C; gas flow: 13 L/min; nebulizer pressure: 60 psig; sheath gas temp: 250°C; sheath gas flow: 12 L/min; VCap: 2000 V; nozzle voltage: 300 V; and fragmentor: 100 V. Data were acquired from 50 to 1700 m/z with active reference mass correction (m/z: 121.0508 and 922.0097) infused through a second nebulizer according to the manufacturer’s instructions, and analyzed in a targeted fashion using the targeted mass spectrometry data analysis software Skyline (v.23.1).

### Purification of recombinant human SUGCT for assays

In order to produce recombinant SUGCT protein for enzyme assays and crystallization, we expressed the most conserved and active isoform 3 (NP_001180242.1, [33]) minus the mitochondrial transit peptide (amino acids 38-445). We initially cloned SUGCT in a Twinstrep-His-SUMO vector, and used this for our enzyme assays. The yield was low, but the protein had activity. Briefly, we inoculated 4L of TB with overnight culture of the his-strep-tagged construct, with 40 μg/ml kanamycin and 1 mM MgSO_4_, grew to OD 2.0, cooled to 16°C and induced overnight with 15 μM IPTG. The next day we resuspended the pellet in TBS with PMSF and lysozyme and lysed using a sonicator (Qsonica). After pelleting down the cellular debris, the lysate was loaded onto a column containing HisPur Ni-NTA Resin (Thermo Scientific) for immobilized metal affinity chromatography (IMAC) purification. The column was equilibrated with 7 column volumes of TBS supplemented with 40 mM imidazole pH 8.0 then washed with 7 column volumes of 20 mM Tris pH 8.0, 50 mM imidazole and adjusted to pH 7.5. The His tagged protein was eluted with 20 mM Tris pH 8.0 containing 150 mM NaCl, 150 mM imidazole and 10% glycerol. We then applied the elution to a streptactin XT column and eluted with streptactin elution buffer (IBA Lifesciences). Then we cleaved the tags with sumo protease overnight. Lastly, we purified the protein on a Superdex200 Increase size exclusion column, and concentrated and froze for assays.

### Purification of SUGCT for crystallography

In order to obtain enough protein for crystallization trials, we screened different epitope tags and found that an N-terminal His tag in a pET47 vector gave the best expression. We expressed the His-SUGCT in Lemo cells (NEB), inoculating 4L of TB with 4 mL overnight culture, 40 μg/ml kanamycin, 1 mM MgSO_4_, and 900 μL of 20% rhamnose. We then grew the cells to OD 2.0, cooled to 16°C and induced overnight with 400 μM IPTG. We then harvested the next day and purified by IMAC the same way as the SUMO construct. After elution, we added HRV3C protease and THP overnight, purified by Superdex200 the next day, concentrated, and flash froze the protein. In order to improve crystallization, we mutated residues Gln 262 and Lys 263 to alanine, which enabled reproducible crystals with losartan carboxylic acid bound. These variants proteins had SUGCT enzyme activity.

### Crystallization and structure determination

We screened crystallization conditions using commercial crystallization screens and identified conditions that gave initial crystals, and then reproduced them with the screening conditions in larger sitting conditions. The final protein crystallized in 0.2 M NaCl, 0.1 M Tris, pH 7.5, 25% PEG 3,350. Crystals appeared after several days, and were flash frozen after transferring to cryoprotectant in liquid nitrogen before collecting at the AMX (17-ID-1) beamline at NSLS II at Brookhaven National Laboratory. Data was processed using autoproc [34] and scaled with Aimless [35]. The structure was solved by molecular replacement, using the alphafold V2 predicted structure as a search model. After molecular replacement, we performed multiple rounds of building using Phenix Autobuild [36]. The models were subsequently refined using Phenix [37] with rigid body refinement and multiple rounds of simulated annealing, minimization, atomic displacement parameter (ADP or B-factor) refinement and TLS refinement (determined using the TLSMD server) [38, 39] with interspersed manual adjustments using Coot [40]. All structural figures were made with PyMOL (The PyMOL Molecular Graphics System, Schrödinger, LLC).

### Recombinant soluble HMG-CoA reductase

The catalytic portion of human HMG-CoA reductase (HMGCR, residues 426–888 [41, 42]) was cloned into the NdeI and BamHI sites of pET-28a using a PCR product generated with the following forward (5’`-`GGC CAT <u>CAT ATG</u> TCA TCA GTA CTG GTG ACA CAG-3’) and reverse primers (5’-AGT ATT <u>GGA TCC</u> TCA GGC TGT CTT CTT GGT GCA AG-3’). Recombinant soluble HMGCR with N-terminal His-tag was produced in *E. coli* and purified using IMAC. The conditions for the HMGCR enzyme assay are: 100 mM potassium phosphate pH7.4, 300 μM NADPH, 150 μM HMG-CoA, 4 mM DTT, 0.1% Triton X-100 and 9 U/L HMGCR. NADPH consumption was followed spectrophotometrically at 340nm using an Agilent/BioTek Synergy H1 plate reader.

### SUGCT enzyme assay

The conditions for the SUGCT enzyme assays are: 100 mM potassium phosphate pH 7.4, 50 μM NADPH, 20 μM glutaryl-CoA, 4 mM DTT, 0.1% Triton X-100 and 9 U/L HMGCR. The enzyme reaction was started with 4 mM HMG unless otherwise noted. NADPH consumption was followed fluorometrically with excitation at 340nm and emission at 460nm using an Agilent/BioTek Synergy H1 plate reader.

## Results

### SUGCT is a modifier of the biochemical phenotype of a GA1 cell line model

We have previously show that in mice, SUGCT deficiency in GA1 mice decreases the accumulation of 3-hydroxyglutaric acid and glutarylcarnitine (C5DC) [12]. In order to establish if SUGCT is able to modify the biochemical phenotype in a human GA1 cell line model, we used CRISPR-Cas9 genome editing to generate *GCDH*/*SUGCT* double KO HEK-293 clonal cell lines. We found that 8 of the 9 *GCDH*/*SUGCT* double KO clones had the anticipated decrease in C5DC when compared to the parental *GCDH* KO clone (**Fig. S2**). Although HEK-293 cells express *SUGCT* mRNA, SUGCT protein was too low to be detected using our validated anti-SUGCT antibody in immunoblotting (**Fig. 1A**). Therefore, the utility of these HEK-293 cell line models to study SUGCT function is limited.

**Figure 1.**
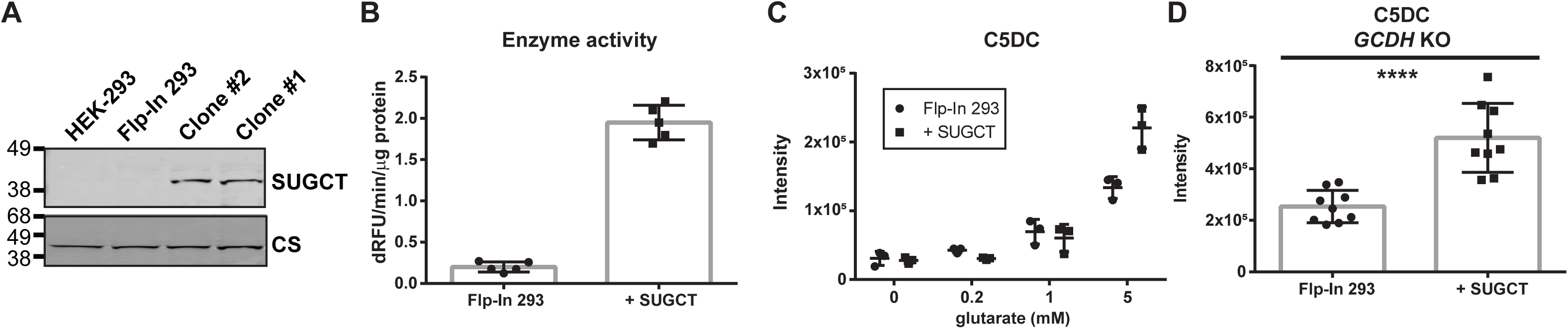
Characterization of SUGCT overexpressing and parental Flp-In 293 cells. (A) Immunoblot showing SUGCT overexpression in 2 representative clones (+ SUGCT). (B) SUGCT enzyme activity. (C) C5DC levels in cell pellets after 24 hours of exposure to different concentration of glutarate in the media. (D) C5DC levels in cell pellets upon GCDH KO using CRISPR-Cas9 genome editing. Three pellets from three independent clones were evaluated. A Mann Whitney test was significant at P < 0.0001.

As an alternative to wild-type HEK-293 cells, we then used the Flp-In system to generate a stable HEK-293 cell line overexpressing high levels of SUGCT. The resulting SUGCT overexpressing cell lines were validated using immunoblot, enzyme and metabolite assays. Whereas SUGCT protein is undetectable in HEK-293 and parental 293 Flp-In cells, it was readily detectable in overexpressing cell lines (**Fig. 1A**). Using our SUGCT assay (described below), we detected increased activity in lysates of overexpressing cell lines (**Fig. 1B**). Importantly, we found that SUGCT overexpressing cell lines accumulate more C5DC upon addition of glutarate to the cell culture media (**Fig. 1C**).

We then used CRISPR-Cas9 genome editing to create a *GCDH* KO in empty Flp-In and SUGCT overexpressing 293 cells as previously described [32, 43, 44]. The SUGCT overexpressing GA1 cell lines had an overall ∼2-fold increase in C5DC, but there is some variability between individual clones (**Fig. 1D**). Variability between CRISPR KO clones has been observed in similar experiments and may be related to off-target effects or heterogeneity of WT cells [45]. We conclude that these HEK-293 cell line models can be used to study the role of SUGCT in GA1.

### A novel fluorometric assay of SUGCT activity

All existing assays of SUGCT activity rely on the separation and quantification of CoA esters using high-performance liquid chromatography (HPLC) with ultra-violet (UV) spectroscopy [2, 3, 33]. In order to enable high throughput screening (HTS) for SUGCT inhibitors, we developed a new spectrophotometric/fluorometric assay to measure SUGCT activity by taking advantage of its known substrate promiscuity. In this assay, SUGCT converts the substrates glutaryl-CoA and HMG into the products glutaric acid and HMG-CoA. The produced HMG-CoA is then measured in a coupled reaction using HMG-CoA reductase. HMG-CoA reductase converts HMG-CoA and 2 NADPH into 2 NADP^+^ and mevalonate (**Fig. 2A**). This conversion of NADPH to NADP^+^ can be monitored spectrophotometrically at 340nm or fluorometrically with excitation at 340nm and emission at 460nm. This new fluorometric assay was optimized for buffer components, as well as enzyme and substrate concentrations. Recombinant SUGCT as measured with this assay displays saturation kinetics for the tested substrates succinyl-CoA (not shown), glutaryl-CoA and HMG (**Fig. S3A**), which are consistent with those reported using an HPLC-based assay [3]. Glutaryl-CoA was used in our assays given its lower cost and likely higher stability as compared to succinyl-CoA. SUGCT accepts a wide variety of dicarboxylic acids as the CoA acceptor, but succinate and glutarate are the preferred substrates. In our assay, dicarboxylic acids are expected to act as competitive inhibitors with respect to HMG, since their conversion to a CoA esters is not detected by HMGCR. Indeed, succinate (C4) and glutarate (C5) were the most effective inhibitors, with malonate (C3), adipate (C6) and suberate (C8) having a higher IC_50_, which closely reflects previous work on SUGCT substrate specificity (**Fig. 2B**) [3, 33]. Indeed, a more detailed kinetic analysis was most consistent with glutarate acting as a competitive inhibitor increasing the apparent K_m_ for HMG (**Fig. S3B**). Combined, these experiments thoroughly validate our novel fluorometric SUGCT assay. Importantly, the results also indicate that the enzyme may be able to cycle between glutarate and glutaryl-CoA.

**Figure 2.**
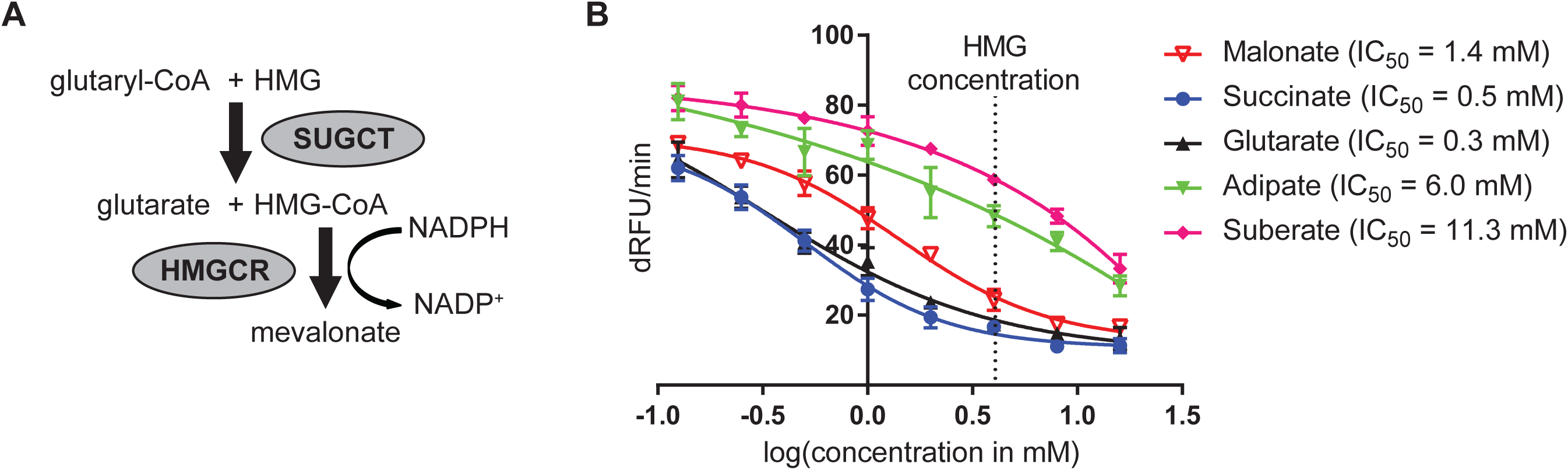
A novel SUGCT enzyme assay. (A) Schema of the SUGCT reaction coupled to HMGCR. (B) IC_50_ curves for malonate, succinate, glutarate, adipate and suberate. HMG concentration is 4 mM.

### High throughput screen of FDA approved library

We then optimized the SUGCT assay to a 384-well plate format and determined the Z-factor, a measure of statistical effect size that serves as an indicator of the quality and performance of the assay in HTS [46]. The Z-factor of our assay was on average 0.53, which is excellent for a HTS. We then screened 853 compounds from an FDA approved drug library. At a screening concentration of 100 μM, we noted a high variation among screening wells. After filtering out wells with likely fluorescent or quenching molecules, the coefficient of variation decreased to 32%. For each well, we then calculated the relative inhibition of SUGCT activity and the number of standard deviations by which the well was above the mean of all screening wells (Z score, **Fig. 3A**). We selected 13 potential hit drugs for repurchase (out of 36 candidates), most with a Z score >3 (minimum 2.7, **Fig. 3A**). Of these, 38% (5) displayed SUGCT inhibition at the screening dose of 100 μM and 15% (2; valsartan and oxaprozin) inhibited at 10 μM (**Fig. 3B**). Valsartan was the only hit with more than 50% inhibition at 10 μM and the IC_50_ was 3.8 μM (**Fig. 3C**). The IC_50_ of oxaprozin was not determined. Of note, we did not identify enzyme activators in this screen (**Fig. 3A**).

**Figure 3.**
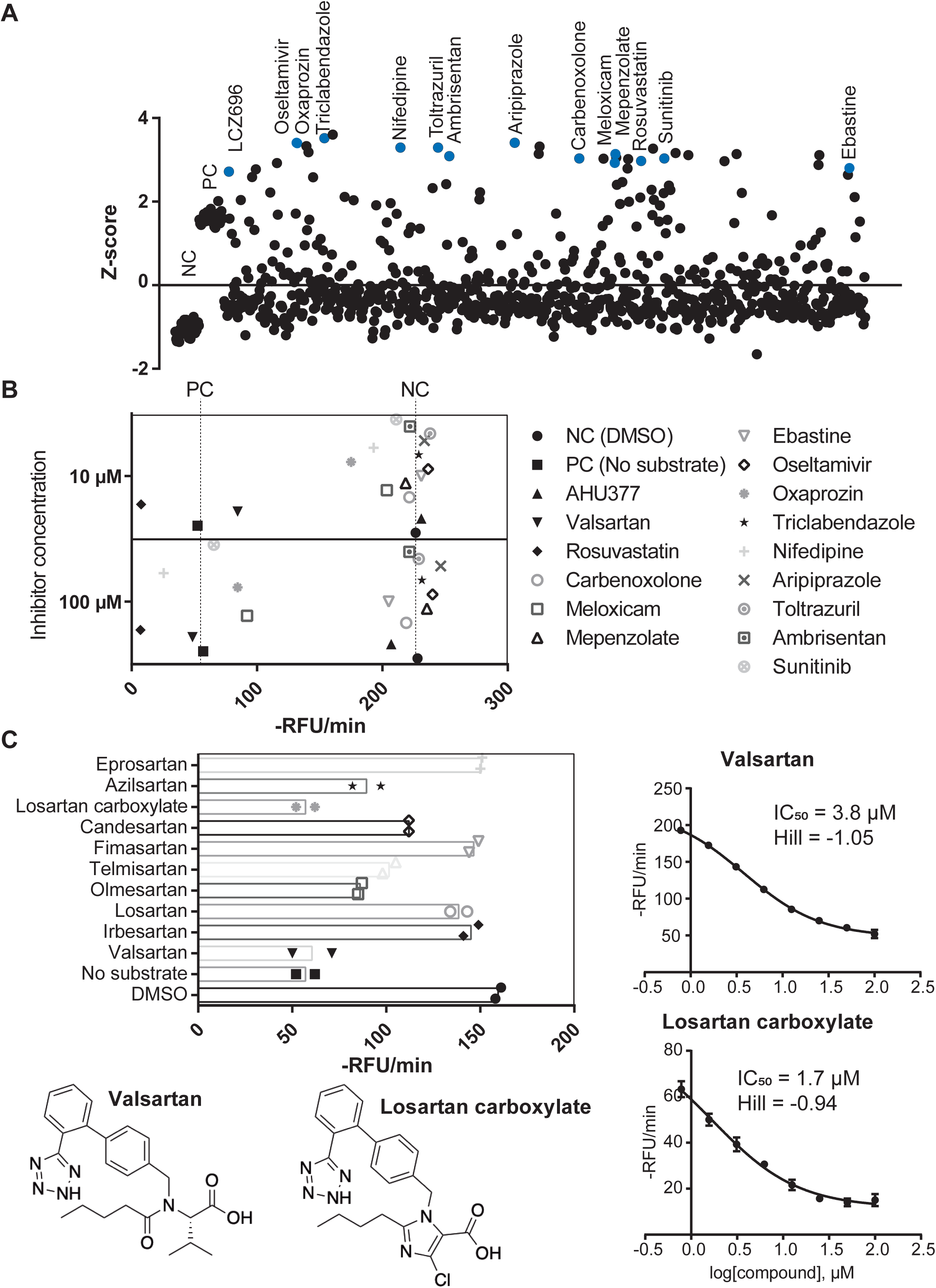
High throughput screen of FDA-approved drugs on SUGCT activity. (A) Z-score of 853 screening wells. Z-scores were calculated for each 384-well plate separately using the average and SD of all screening wells after removal of outliers due to compound fluorescence or quenching. Negative control (NC, DMSO) and positive control (PC, no enzyme) not included in the calculation of the average and SD. Compounds selected for repurchase are indicated in blue. (B) Repurchase of 15 FDA approved drugs. Each compound was measured in duplicate at 100 µM and 10 µM. The hit LCZ969 is a fixed-dose combination medication containing Sacubitril (AHU377) and valsartan. Rosuvastatin was selected as positive control. Therefore 5/13 hits (38%) confirmed at the 100 µM screening dose. (C) Effect of 10 different Angiotensin II Receptor Blockers (ARBs) on SUGCT activity. Each ARB was measured in duplicate at 10 µM. Telmisartan displayed some fluorescence, which may explain the lower RFU/min. (D) IC_50_ curve for valsartan. (E) IC_50_ curve for losartan carboxylic acid.

Valsartan belongs to the drug class of Angiotensin II Receptor Blockers (ARBs) that are widely used to treat high blood pressure and heart failure. Subsequently, we purchased and evaluated 9 additional related ARBs (**Fig. 3C**). The carboxylic acid metabolite of losartan (E-3174) was slightly more active than valsartan (IC_50_ = 1.7 μM), while the hydroxy parent drug losartan was inactive, revealing that the carboxylate group is essential for inhibitory activity (**Fig. 3C**). The ethyl ester of valsartan also had an IC_50_ that was 40 times larger, further demonstrating the importance of a free carboxylate. All other ARBs had either low (olmesartan and azilsartan) or no activity. This pilot HTS identified the first hit SUGCT inhibitors with some preliminary structure-activity relationships (SAR) demonstrating that the ARB core (only valsartan and losartan carboxylic acid) and the carboxylic acid group are important for SUGCT inhibition.

The presence of a free carboxylate in valsartan and losartan carboxylic acid could suggest that these ARBs are competitive inhibitors with respect to HMG and could be converted into a CoA ester. We found no evidence that SUGCT was able to convert valsartan into a CoA ester (**Fig. S4**).

Our SUGCT assay is indirect and uses HMGCR as auxiliary enzyme. The assay is therefore also expected to identify HMGCR inhibitors. Indeed, one of our hits was rosuvastatin (**Fig. 3A**), an HMGCR inhibitor. Therefore, valsartan and losartan carboxylic acid were counter-screened in a spectrophotometric HMGCR assay and did not inhibit up to 100 μM, confirming that they are true SUGCT inhibitors without activity on HMGCR which would result in a false positive hit result.

### The crystal structure of the human SUGCT

After optimizing conditions to purify large quantities of SUGCT, we undertook crystallization trials, in order to determine the structure of the protein. Using our purified protein, we set up crystallization trials with valsartan and were able to obtain crystals that diffracted to 2.4 Å. We then solved the structure by molecular replacement using the AlphaFold V2 structure as a search model (Figure 4 and Table S1). Based on the gel filtration (**Fig. S5**), it was expected to find a monomer. However, molecular replacement was able to place two copies that form an interlocking dimer, explaining the small hydrodynamic radius. The active site is formed at the interface of the two monomers, indicated by the conserved catalytic Asp212 residue found on each side of the dimer (**Fig. 4A**). The monomers are interlocking, as seen in the surface view (**Fig. 4B**), suggesting they can only fold correctly as a dimer which was not predicted by Alphafold. No density for valsartan bound was detected, likely due to solubility problems, so this structure was deemed an apo structure.

**Figure 4.**
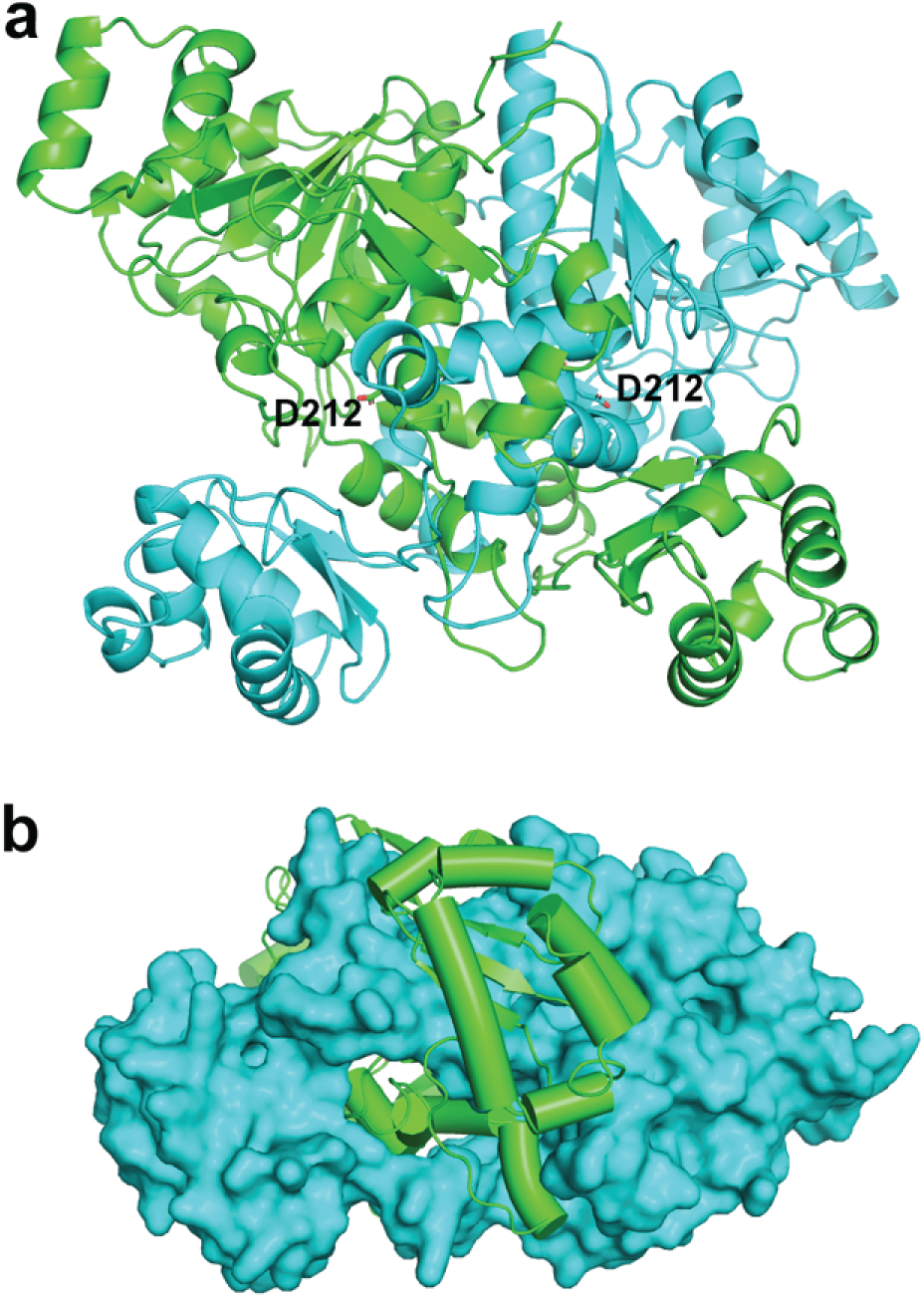
Overall structure of SUGCT. (A) Dimeric structure with catalytic residue (Asp212) shown. (B) Surface and cartoon view of SUGCT dimer, indicating dimeric assembly of polypeptide chains.

The SUGCT structure is highly conserved from prokaryotic type-III CoA transferase, like the crotonobetainyl-CoA: carnitine CoA-transferase (CaiB) from *E. coli* [5, 6], the formyl-CoA transferase from *Oxalobacter formigenes* [9, 10] and *E. coli* [7, 8], or the α-methylacyl-CoA racemase from *Mycobacterium tuberculosis* [11]. Despite only low sequence identity (**Fig. S6**), the structures are similar, with an RMSD of 1.7 Å^2^. The large and open active site formed at this interface can accommodate the large substrates and can help explain the promiscuity of the enzyme for different substrates. Consistent with previous studies on these homologues, Asp212 in SUGCT is conserved and positioned in the active site, consistent with being a catalytic residue. We found that the p.Asp212Ala variant folded properly, but was inactive consistent with a crucial role in catalysis (**Fig. S7**).

The majority of the GA3 cases are caused by the p.Arg336Trp variant. The high allele frequency make this variant (rs137852860) of particular in interest. Although this residue is not conserved, earlier work demonstrated that the variant protein is much less stable than the wild-type protein [33]. We used our SUGCT structure to better understand the consequences of this amino acid substitution on the SUGCT structure. Arg 336 forms a salt bridge with Glu 339 and a hydrogen bond with the carbonyl backbone of Leu 305, which are key interactions for folding (**Fig. S8A**). The p.Arg322Trp variant, homozygous in one GA3 case [22], also has a significant number of carriers in gnomAD. Functional studies for this variant have not been reported. Arg 322 makes two important hydrogen bonds, one with the carbonyl backbone of Val 276 and another with the carbonyl backbone of Tyr 316 (**Fig. S8B**).

### The crystal structure of the human SUGCT in complex with losartan carboxylic acid

In order to understand inhibition of SUGCT, we then attempted to crystallize SUGCT with losartan carboxylic acid, which was the most potent inhibitor from our pilot screen. After crystallization trials with a SUGCT variant designed to have improved crystal packing [47], we obtained a new crystal form of SUGCT that contained two dimers. There was unambiguous density for losartan carboxylic acid in one of the four active sites (**Fig. S9**), and density consistent with the inhibitor in two of the other active sites. In the fourth site, predominantly from chain B, we observed some unmodeled density that was not consistent with losartan. It is possible this density was from some residual substrate from purification in *E. coli* that remained bound in this condition, but it was not clear enough to model in CoA or other derivatives.

Losartan carboxylic acid binds in the active site, near where glutaric acid likely binds (**Fig. 5A**). The inhibitor binds in a pocket formed by residues from both chains of the dimer (**Fig. 5B**). Interestingly, the inhibitor buries deep into a novel pocket that does not exist in our apo structure. This new pocket is caused by large rotations of the side chains mostly from three residues (Trp91, Tyr102, and His253), compared to the apo structure (**Fig. 5C**).

**Figure 5.**
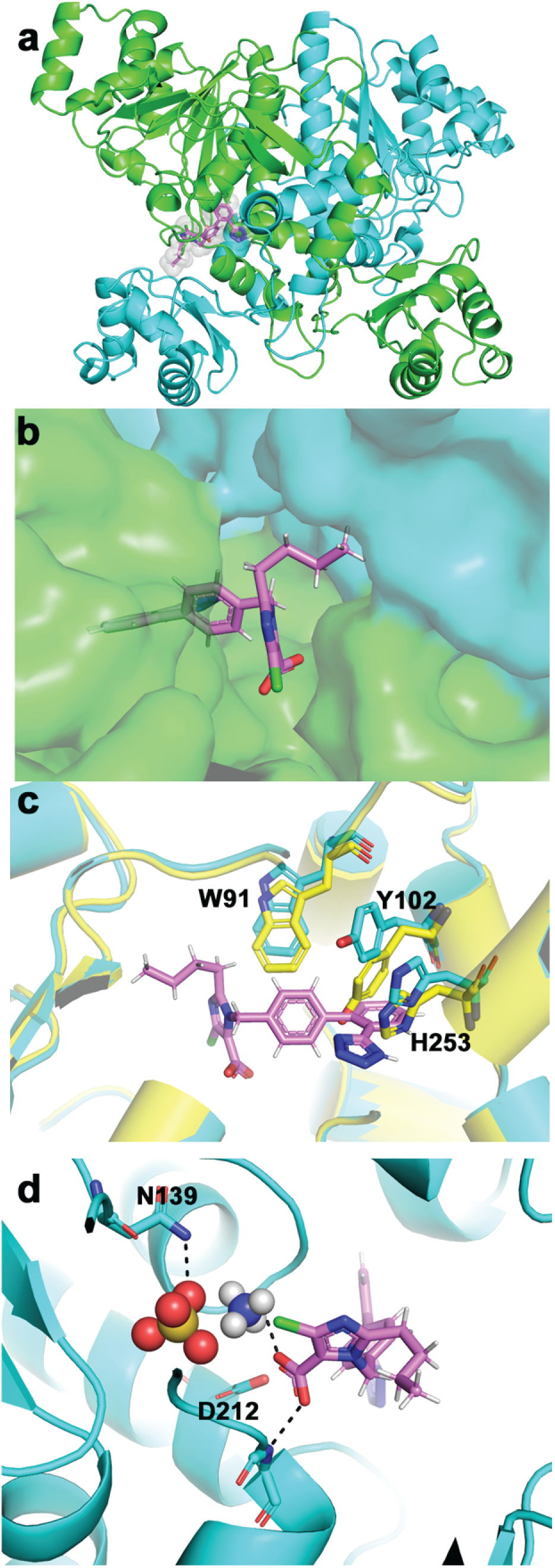
Structure of SUGCT:Losartan carboxylic acid complex. (A) Overall structure of the complex with Losartan carboxylic shown in purple in the left active site. (B) Close-up view of Losartan bound in the pocket, colored by chain. (C) Comparison of apo (yellow) and Losartan (cyan) active sites. Key residues are shown in stick form. (D) Close-up view of the losartan carboxylic acid interaction with the catalytic site. Ammonium (blue and white) and sulfate (red and orange) ions are shown in sphere form, with potential interactions shown with dashed lines.

One perplexing aspect of the structure is the placement of the carboxylic acid of losartan in close proximity of the catalytic Asp212. We then noticed additional density in the active site, corresponding to two spheres. We modeled these as ammonium and sulfate ions, which were present in the crystallization conditions (**Fig. 5D**). Indeed, after extensive screening, we were only able to obtain crystals with ammonium sulfate or ammonium acetate salts, suggesting that they were playing a role in binding the losartan. The ammonium ion neutralizes the losartan carboxylic acid, which is also stabilized by the unsatisfied amide backbone of the helix and the helix dipole, which we have previously observed to stabilize phosphates [48]. We think therefore, that there is a favorable interaction with the losartan carboxylic acid and the N-terminus of the helix, which explains the preference for the carboxylic acid form of losartan over losartan itself, with hydroxyl group, for inhibition of SUGCT. Moreover, the structure suggested that the salt is key for inhibition in vitro. In our assay buffer, we had potassium phosphate, which has a similar size to the ammonium sulfate in our crystallization condition. Therefore, we decided to test different buffers to see the effect on the IC_50_. By altering both the cation and anions, we were able to study whether this model of the salt binding is important for inhibition. Indeed, we saw an increase in IC_50_ if we removed either the phosphate or the potassium from the reaction mixture (**Fig. S10**). Most notable was a 5.5-fold increase in the IC_50_ upon switching from phosphate to Hepes buffer (**Fig. S10**). This indicates that the cations and anions in the carboxylic acid binding site are important for inhibition in the assay. Although potassium and phosphate are physiological ions in the cell, it is important to note that the buffer identity can play a role in the inhibition, which will inform future inhibition studies and inhibitor designs.

## Discussion

SUGCT is an auxiliary enzyme in lysine degradation. This type-III CoA transferase catalyzes the succinyl-CoA-dependent re-esterification of free glutarate back into glutaryl-CoA. This reaction is a metabolite repair pathway, necessary to prevent urinary loss of glutarate. Mouse studies have shown that SUGCT is a modifier of the biochemical phenotype of GA1 [12]. SUGCT deficiency decreases the accumulation of potentially neurotoxic metabolites including 3-hydroxyglutaric acid [12]. Herein, we developed chemical and biological tools useful for the future discovery of a high affinity inhibitor of SUGCT. Using heterologous expression of SUGCT, we were able to solve the first crystal structure of human SUGCT at 2.4 Å resolution. We also developed a SUGCT enzyme assay which we demonstrate is amenable for an HTS of larger chemical libraries. This assay was validated in a pilot screen of an FDA approved drug library that identified valsartan as a dose dependent SUGCT inhibitor with low micromolar IC_50_. Losartan carboxylic acid, the active metabolite of losartan, was subsequently identified as an analog with similar ability to inhibit SUGCT. We also developed HEK-293 cell line models that can be used to validate in cell activity of SUGCT inhibitors. Unfortunately, we did not observe a decrease in C5DC upon treatment of SUGCT overexpressing GA1 HEK-293 cells with the parent drug or the presumably more permeable valsartan ethyl ester. Additional drug development including a larger HTS is necessary to further pharmacologically validate SUGCT as a small molecule drug target to treat GA1.

It is notable that SUGCT is able to use glutarate and glutaryl-CoA as substrates and thus essentially performing a zero-sum reaction. In GA1 disease state, wherein there are elevated levels of both glutaryl-CoA and glutaric acid, this reaction may constitute a significant portion of the gross SUGCT enzyme flux. However, our previous study of GA1 mice with defective SUGCT has demonstrated that even in this condition with extreme elevation of glutaryl-CoA, the net flux through SUGCT leads to resynthesis of glutaryl-CoA [12]. We speculate that this is enabled by readily available succinyl-CoA for example as a result of substrate channeling. Indeed, succinyl-CoA and glutaryl-CoA are produced by two separate enzymes, the 2-oxoglutarate dehydrogenase complex (OGDHc) and 2-oxoadipate dehydrogenase complex, respectively. We therefore speculate that SUGCT is in proximity to OGDHc.

We solved the first structure of a eukaroytic type-III CoA transferase, which highlights the conservation of this family. We have also identified valsartan and losartan carboxylic acid as inhibitors of SUGCT. These ARBs, however, are unlikely to inhibit SUGCT *in vivo* given their low potency and poor cell permeability. Indeed, we were unable to detect activity in our SUGCT overexpressing GA1 HEK-293 cell line. Moreover, we did not find any literature reports of increased glutaric acid in urine of people treated with valsartan or losartan. Urinary organic acid analysis is, however, not a standard clinical chemistry test, and therefore such an analysis may be worthwhile to pursue.

Valsartan and losartan carboxylic acid may not be useful drugs for targeting SUGCT. However, the identification of preliminary inhibitors hits and determination of the co-crystal structure in complex with SUGCT provide several important insights. First, our screen is validated by the discovery of inhibitors, which will facilitate a future HTS to find novel hit compounds from a larger unbiased library. Secondly, the structure revealed a novel binding mode that induces a new pocket not previously visible for this family of enzymes. The new structure therefore may facilitate inhibitor design targeting this pocket.

## Supporting information

All supplemental material

## Acknowledgements

Research reported in this publication was supported by the Eunice Kennedy Shriver National Institute of Child Health & Human Development and the National Institute of General Medical Sciences of the National Institutes of Health under Award Numbers R21HD102745 (to S.M.H. and R.J.D.) and R35GM124838 (to M.B.L.). This research used resources of the National Synchrotron Light Source II, a US Department of Energy (DOE) Office of Science User Facility operated for the DOE Office of Science by Brookhaven National Laboratory, under Contract No. DESC0012704. The Life Science Biomedical Technology Research resource is primarily supported by the NIH, National Institute of General Medical Sciences (NIGMS) through a Biomedical Technology Research Resource P41 grant (No. P41GM111244), and by the DOE Office of Biological and Environmental Research (No. KP1605010). This work was supported in part through the computational and data resources and staff expertise provided by Scientific Computing and Data at the Icahn School of Medicine at Mount Sinai and supported by the Clinical and Translational Science Award (CTSA) grant UL1TR004419 from the National Center for Advancing Translational Sciences.

