## Supplementary material for "Characterization, structure and inhibition of the human succinyl-CoA:glutarate-CoA transferase, a genetic modifier of glutaric aciduria type 1": All supplemental material

Supplementary Tables and Supplementary Figures

#### Supplementary Tables

Table S1. Crystallography collection and processing statistics.

| | SUGCT apo | SUGCT-Losartan complex |
|---|---|---|
| **Data collection** | | |
| Space group | C 1 2 1 | P212121 |
| Cell dimensions | | |
| *a*, *b*, *c* (Å) | 91.08 115.13 79.22 | 107.93 119.81 136.25 |
| α, β, γ (°) | 90.00 100.79 90.00 | 90.00 90.00 90.00 |
| Resolution (Å) | 48.84 - 2.40 (2.49 - 2.40)* | 34.31 - 2.08 (2.12 - 2.08) |
| $R_{sym}$ or $R_{merge}$ | 0.094 (1.368) | 0.122 (1.694) |
| $I / \sigma I$ | 7.0 (0.8) | 6.8 (0.9) |
| Completeness (%) | 99.8 (99.8) | 99.69 (99.92) |
| Redundancy | 3.5 (3.2) | 4.8 (5.0) |
| $CC_{1/2}$ | 0.996 (0.406) | 0.997 (0.420) |
| **Refinement** | | |
| Resolution (Å) | 44.74 - 2.4 (2.486 - 2.4) | 33.41 - 2.08 (2.154 - 2.08) |
| Total reflections | 110117 (9838) | 106146 (10540) |
| $R_{work}$ / $R_{free}$ | 0.2201 / 0.2389 | 0.2220 / 0.2483 |
| No. atoms | 6219 | 12726 |
| Protein | 6214 | 12378 |
| Ligand/ion | 0 | 199 |
| Water | 5 | 218 |
| *B*-factors | | |
| Protein | 68.49 | 52.44 |
| Ligand/ion | | 52.42 |
| Water | 63.91 | 44.55 |
| R.m.s. deviations | | |
| Bond lengths (Å) | 0.003 | 0.002 |
| Bond angles (°) | 0.53 | 0.494 |

* Data is from a single crystal. Parentheses indicate highest resolution shell.

#### Supplementary Figure legends

**Supplementary Figure 1. Overview of SUGCT function.** (A) SUGCT catalyzes the succinyl-CoA-dependent conversion of glutaric acid into succinate and glutaryl-CoA (EC 2.8.3.13). (B) Metabolism of glutaryl-CoA in GA1 patients. GCDH is the enzyme deficient in GA1. Formation of glutaric acid from glutaryl-CoA may be non-enzymatic or mediated by a thioesterase or another enzyme [14, 15]. C5DC is formed by transesterification of glutaryl-CoA with free carnitine. The red arrows indicate the direction of change in the indicated metabolite in GA1 mice with an additional defect in SUGCT (see below [12]).

**Supplementary Figure 2. Characterization of *GCDH*/*SUGCT* double KO cell lines.** One *GCDH* KO clone (G9 [44]) was selected for the generation of the *GCDH*/*SUGCT* double KO cell lines. Analysis of glutarylcarnitine (C5DC) in the *GCDH*/*SUGCT* double KO cell lines, the parental *GCDH* single KO cell line and wild-type HEK-293 cells. Single *GCDH* KO clones are indicated as G# and double *GCDH*/*SUGCT* KO clones as G#S#. Three *GCDH*/*SUGCT* KO were confirmed through Sanger sequencing of the targeted *SUGCT* exon.

**Supplementary Figure 3. SUGCT enzyme kinetics.** (A) SUGCT enzyme in RFU/min activity versus substrate concentration. Left panel: varying glutaryl-CoA with 5 mM HMG. Right panel: varying HMG with 40 μM glutaryl-CoA. (B) SUGCT enzyme in RFU/min activity versus substrate concentration in the presence of varying concentrations of the inhibitor glutarate. Left panel: varying glutaryl-CoA with 6.7 mM HMG. Right panel: varying HMG with 40 μM glutaryl-CoA.

**Supplementary Figure 4. Valsartan and CoA transfer.** HMG-CoA was incubated with SUGCT and the compounds listed in the graphs. If any of those compounds is a SUGCT substrate, the HMG-CoA will be converted into the respective other CoA. After 30 minutes, NADPH and HMGCR were added to convert the remaining HMG-CoA into $NADP^+$ and mevalonate. RFU indicates the level of NADPH at the endpoint of the assay, with higher values representing lower HMG-CoA available to convert the fluorescent NADPH into non-fluorescent $NADP^+$. Glutarate and succinate are converted to CoA esters, but not valsartan. As demonstrated before, valsartan inhibits conversion of glutarate and succinate.

**Supplementary Figure 5. FPLC trace of SUGCT.** Curve is shown for purification on Superdex200 increase.

**Supplementary Figure 6. Alignment of different type-III CoA transferases.** CLUSTAL O (1.2.4) multiple sequence alignment. We used the most recent SUGCT protein sequence (NP_001180242.2) for this alignment. This sequence misses the 7 N-terminal amino acids (MPSETHA) that were included in the original sequence (NP_001180242.1) as part of the mitochondrial targeting sequence. Throughout the manuscript we have used the amino acid numbering according to NP_001180242.1. Selected residues in SUGCT are indicated. The amino acid number according to NP_001180242.2 is indicated in parentheses.

**Supplementary Figure 7. Activity of the p.D212A SUGCT variant.** Enzyme activity of two batches of wild-type SUGCT and the p.D212A variant were measured at different protein concentrations. One representative mid-range concentration is shown. Even at the highest protein concentration, the p.D212A variant had no detectable enzyme activity.

**Supplementary Figure 8. SUGCT residues observed as variants in individuals with GA3.** (A) Arg336 residue mutated as p.R336W. (B) Arg322 residue mutated as p.R322W.

**Supplementary Figure 9. Electron density for losartan carboxylic acid.** An Fo-Fc map was generated by omitting the inhibitor and refining in Phenix. The map is shown at 2 sigma contour.

**Supplementary Figure 10. Valsartan $IC_{50}$ curves and buffer effects.** $IC_{50}$ curves for valsartan using different biological buffer systems. The standard assay uses 100 mM potassium phosphate (KPi) buffer pH 7.4 ($IC_{50}$ = 1.944). The other conditions are 100 mM

Hepes buffer adjusted with KOH to pH 7.4 ($IC_{50}$ = 10.67), 100 mM Hepes buffer adjusted with NaOH to pH 7.5 ($IC_{50}$ = 18.04), 100 mM Tris buffer adjusted with HCl to pH 7.44 ($IC_{50}$ = 16.30) and 100 mM Tris buffer adjusted with HCl to pH 7.44 with 50 mM KCl ($IC_{50}$ = 12.12). Same amounts of enzyme used for each condition indicate effects on enzyme activity.

#### Figure S1

##### A. SUGCT enzyme reaction

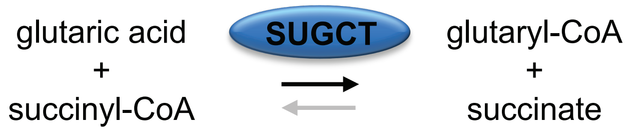


##### B. SUGCT inhibition in GA1

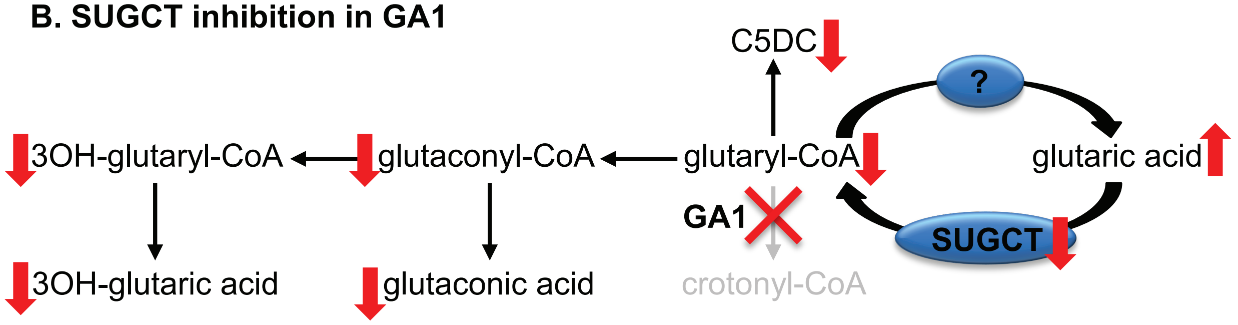

#### Figure S2

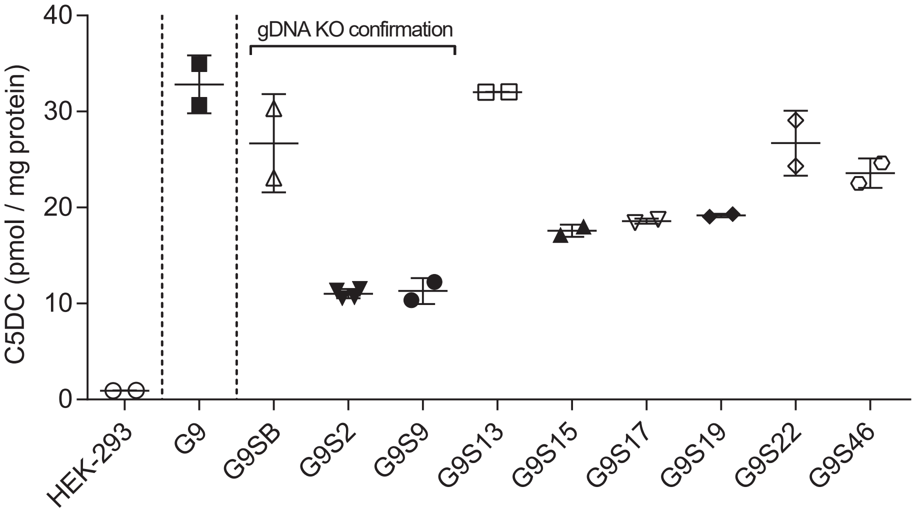
gDNA KO confirmation
C5DC (pmol / mg protein)
40
30
20
10
0
HEK-293
G9
G9SB
G9S2
G9S9
G9S13
G9S15
G9S17
G9S19
G9S22
G9S46

**Figure S3**

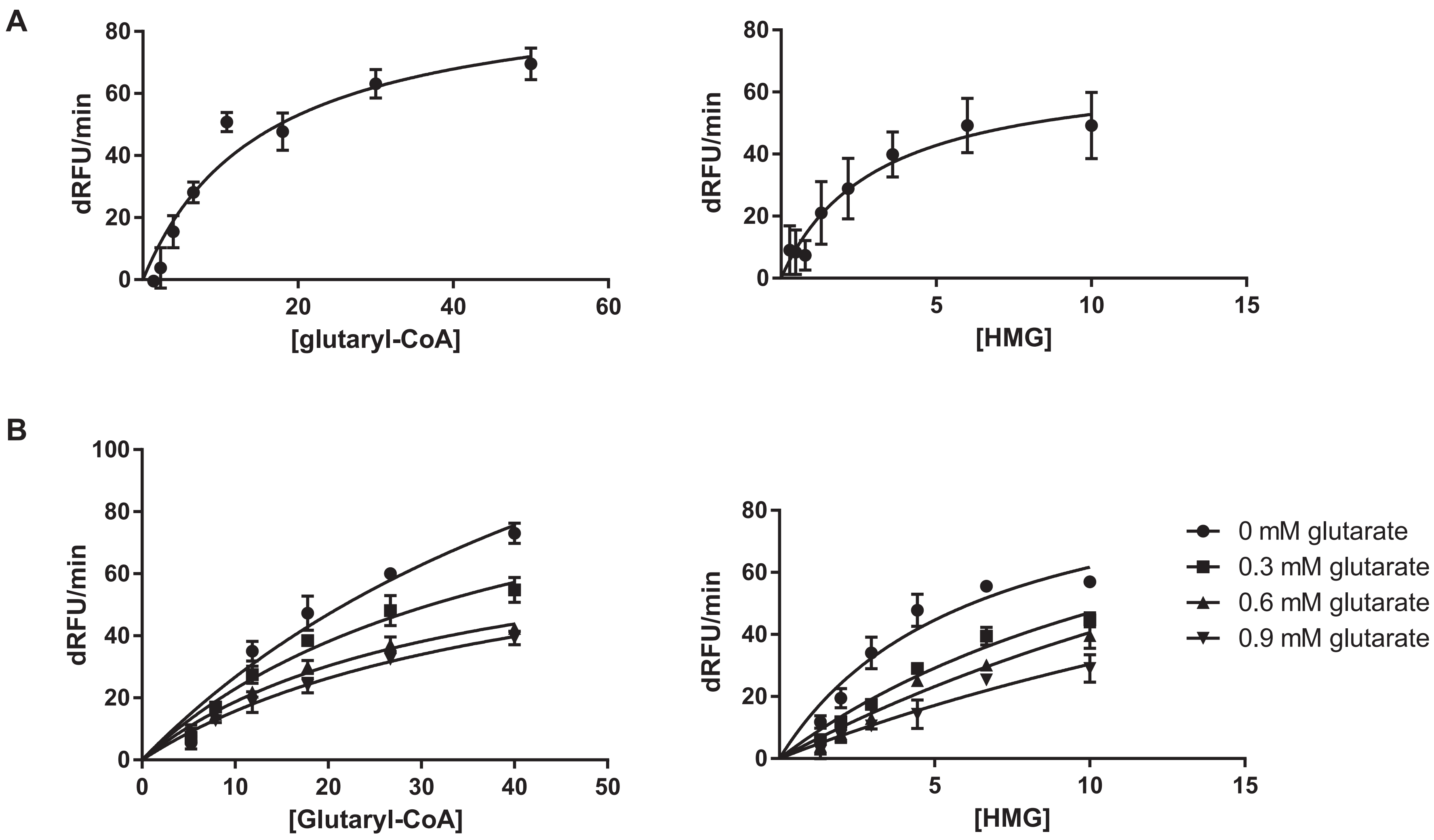

A
dRFU/min
[glutaryl-CoA]
[HMG]
B
[Glutaryl-CoA]
0 mM glutarate
0.3 mM glutarate
0.6 mM glutarate
0.9 mM glutarate

**Figure S4**

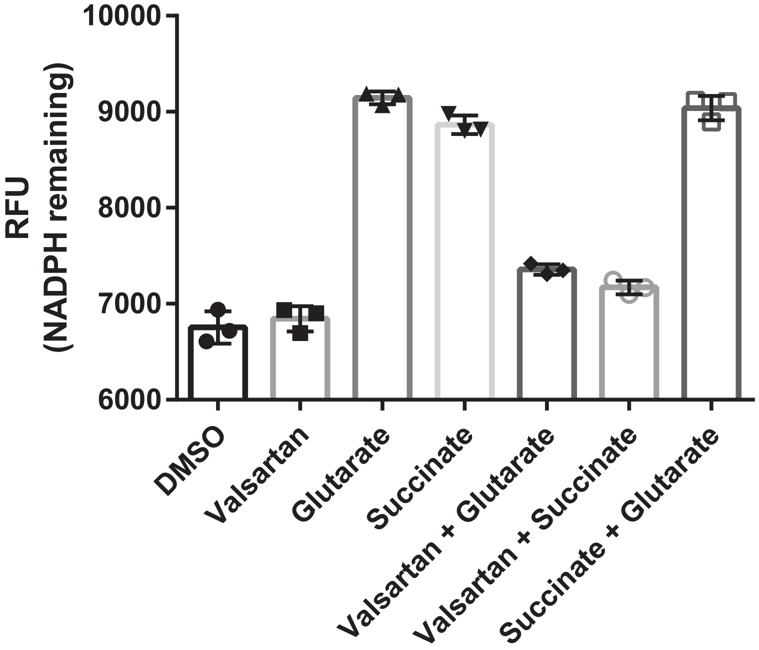

RFU
(NADPH remaining)
10000
9000
8000
7000
6000
DMSO
Valsartan
Glutarate
Succinate
Valsartan + Glutarate
Valsartan + Succinate
Succinate + Glutarate


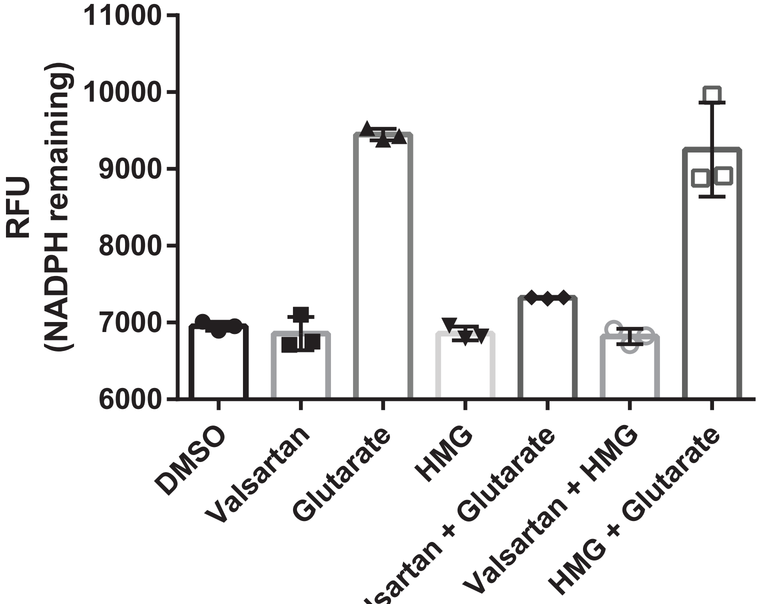

RFU
(NADPH remaining)
11000
10000
9000
8000
7000
6000
DMSO
Valsartan
Glutarate
HMG
Valsartan + Glutarate
Valsartan + HMG
HMG + Glutarate

#### Figure S5

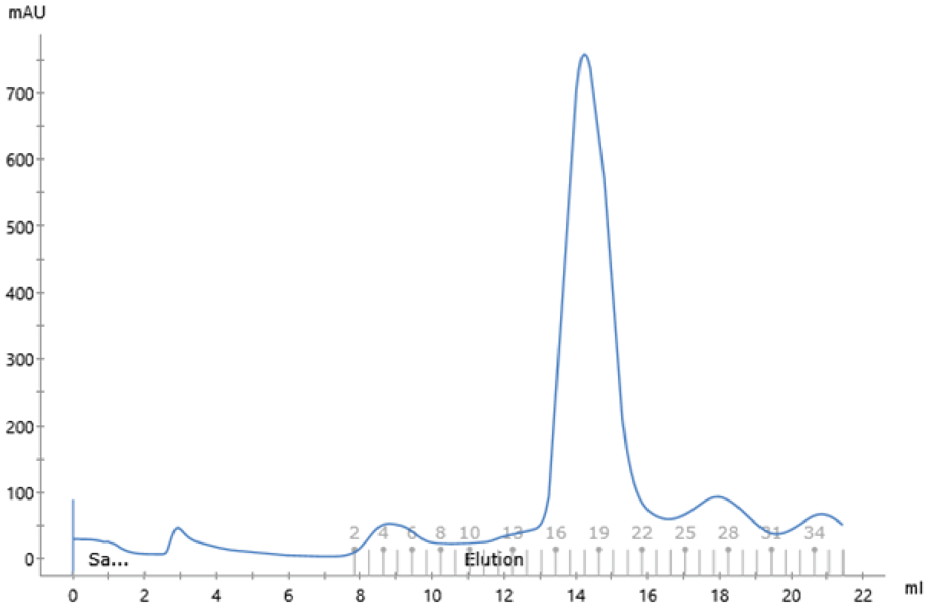

mAU
700
600
500
400
300
200
100
0
Sa...
Elution
2 4 6 8 10 13 16 19 22 25 28 31 34
0 2 4 6 8 10 12 14 16 18 20 22
ml

### Figure S6

```
CLUSTAL O(1.2.4) multiple sequence alignment
```

Mitochondrial targeting sequence

```
NP_001180242.2             MLATLARVAALRRTCLFSGRGGGRGLWTGRPQSDMNNIKPLEGVKILDLTRVLAGPFATM 60
sp|O06644|FCTA_OXAFO       ------------------------------------MTKPLDGINVLDFTHVQAGPACTQ 24
sp|P69902|FCTA_ECOLI       ------------------------------------MSTPLQGIKVLDFTGVQSGPSCTQ 24
sp|O06543|AMACR_MYCTU      ------------------------------------MAGPLSGLRVVELAGIGPGPHAAM 24
sp|P31572|CAIB_ECOLI       -----------------------------MDHLPMPKFGPLAGLRVVFSGIEIAGPFAGQ 31
                                                                  ** *:.::       ** .

NP_001180242.2             NLGDLGAEVIKVERPGAGDDTRTWGPPFVGTESTYYLSVNRNKKSIAVNIKDPKGVKIIK 120
sp|O06644|FCTA_OXAFO       MMGFLGANVIKIERRGSGDMTRGWLQDKPNVDSLYFTMFNCNKRSIELDMKTPEGKELLE 84
sp|P69902|FCTA_ECOLI       MLAWFGADVIKIERPGVGDVTRHQLRDIPDIDALYFTMLNSNKRSIELNTKTAEGKEVME 84
sp|O06543|AMACR_MYCTU      ILGDLGADVVRIDRPSSVDGIS--------RDA-----MLRNRRIVTADLKSDQGLELAL 71
sp|P31572|CAIB_ECOLI       MFAEWGAEVIWIENVAWADTIR--------VQPNYPQLSRRNLHALSLNIFKDEGREAFL 83
                            :.  **:*: ::. .  *            :         * : :  :    :* :

NP_001180242.2             ELAAVCDVFVENYVPGKLSAMGLGYEDIDEIAPHIIYCSITGYGQTGPISQ--RAGYDAV 178
sp|O06644|FCTA_OXAFO       QMIKKADVMVENFGPGALDRMGFTWEYIQELNPRVILASVKGYAEGHANEH--LKVYENV 142
sp|P69902|FCTA_ECOLI       KLIREADILVENFHPGAIDHMGFTWEHIQEINPRLIFGSIKGFDECSPYVN--VKAYENV 142
sp|O06543|AMACR_MYCTU      KLIAKADVLIEGYRPGVTERLGLGPEECAKVNDRLIYARMTGWGQTGPRSQQAGHDINYI 131
sp|P31572|CAIB_ECOLI       KLMETTDIFIEASKGPAFARRGITDEVLWQHNPKLVIAHLSGFGQYGTEEYTNLPAYNTI 143
                           ::    *:::*          *:  *   :   :::   :.*: :            : :

                                                     ↓
NP_001180242.2             ASAVSGLMHITGPENGDPVRPGVAMTDLATGLYA-YGAIMAGLIQKYKTGKGLFIDCNLL 237
sp|O06644|FCTA_OXAFO       AQCSGGAAATTGFWDGPPTVSGAALGDSNSGMHL-MIGILAALEMRHKTGRGQKVAVAMQ 201
sp|P69902|FCTA_ECOLI       AQAAGGAASTTGFWDGPPLVSAAALGDSNTGMHL-LIGLLAALLHREKTGRGQRVTMSMQ 201
sp|O06543|AMACR_MYCTU      S--LNGILHAIGRGDERPVPPLNLVGDFGGGSMFLLVGILAALWERQSSGKGQVVDAAMV 189
sp|P31572|CAIB_ECOLI       AQAFSGYLIQNG-DVDQPMPAFPYTADYFSGLTA-TTAALAALHKVRETGKGESIDIAMY 201
                           :   .*     *     *        *   *      . :*.*    .:*:*  :   :
```

Catalytic residue bold underlined Asp212 (205) in SUGCT

```
NP_001180242.2             SSQVACL-S-----------HIAANYLIGQ-----------------KEAKRWGTA---- 264
sp|O06644|FCTA_OXAFO       DAVLNLVRIKLRDQQRLERTGILAEYPQAQPNFAFDRDGNPLSFDNITSVPRGGNAGGGG 261
sp|P69902|FCTA_ECOLI       DAVLNLCRVKLRDQQRLDKLGYLEEYPQYPNGT-F-----------GDAVPRGGNAGGGG 249
sp|O06543|AMACR_MYCTU      DGSSVLIQMMWA---------MRATG--------M------------WTDTRGANMLDGG 220
sp|P31572|CAIB_ECOLI       EVMLRMGQYFM----------M--DY--------F------------NGGEMCPRMSKGK 229
                           .

                                                                                    ↓
NP_001180242.2             -HGSIVPYQAFKTKDGYIVVGAGNNQQFATVCKILDLPEL-IDNSKYKTNH-----LRVH 317
sp|O06644|FCTA_OXAFO       QPGWMLKCKGWETDADSYVYFTIAANMWPQICDMIDKPEW-KDDPAYNTFE-----GRVD 315
sp|P69902|FCTA_ECOLI       QPGWILKCKGWETDPNAYIYFTIQEQNWENTCKAIGKPEW-ITDPAYSTAH-----ARQP 303
sp|O06543|AMACR_MYCTU      -APYYD---TYECADGRYVAV----GAIEPQFYAAMLAGLGLDAAELPPQNDRA---RW- 268
sp|P31572|CAIB_ECOLI       -DPYYAGCGLYKCADGYIVMELVGITQIEECFKDIGLAHL-LGTPEIPEGTQLIHRIECP 287
                                     ::   .  :                                      .
```

Arg322 (315) bold underlined

```
                                      ↓
NP_001180242.2             NRKELIKILSERFEEELTSKWLYLFEGSGVPYGPINNMKNVFAEPQVLHNGLVMEMEH-- 375
sp|O06644|FCTA_OXAFO       KLMDIFSFIETKFADKDKFEVTEWAAQYGIPCGPVMSMKELAHDPSLQKVGTVVEVVD-- 373
sp|P69902|FCTA_ECOLI       HIFDIFAEIEKYTVTIDKHEAVAYLTQFDIPCAPVLSMKEISLDPSLRQSGSVVEVEQ-- 361
sp|O06543|AMACR_MYCTU      --PELRALLTEAFASHDRDHWGAVFANSDACVTPVLAFGEVHNEPHIIERNTFYEANGGW 326
sp|P31572|CAIB_ECOLI       YGPLVEEKLDAWLATHTIAEVKERFAELNIACAKVLTVPELESNPQYVARES----ITQW 343
                               :   :          .        .     :  . ::  :*
```

Arg336 (329) bold underlined

```
NP_001180242.2             -PTVGKI-SVPGPAVRYSKFKMSEARPPPLLGQHTTHILKEVLRYDDRAIGELLSAGVVD 433
sp|O06644|FCTA_OXAFO       -EIRGNH-LTVGAPFKFSGFQPE-ITRAPLLGEHTDEVLKEL-GLDDAKIKELHAKQVV- 428
sp|P69902|FCTA_ECOLI       -PLRGKY-LTVGCPMKFSAFTPD-IKAAPLLGEHTAAVLQEL-GYSDDEIAAMKQNHAI- 416
sp|O06543|AMACR_MYCTU      QPMPAP---------RFSRTASSQ-PRPPAATIDIEAVLTDW-DG--------------- 360
sp|P31572|CAIB_ECOLI       QTMDGRTCKGPNIMPKFKNNPGQIWRGMPSHGMDTAAILKNI-GYSENDIQELVSKGLAK 402
                               .          ::.    .     *    .   :* :

NP_001180242.2             QHETH 438
sp|O06644|FCTA_OXAFO       ----- 428
sp|P69902|FCTA_ECOLI       ----- 416
sp|O06543|AMACR_MYCTU      ----- 360
sp|P31572|CAIB_ECOLI       VED-- 405
```

### Figure S7

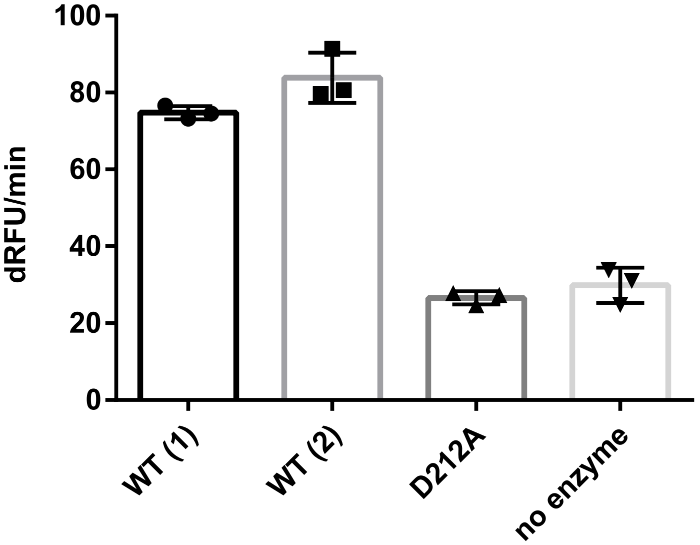

**Figure S8**

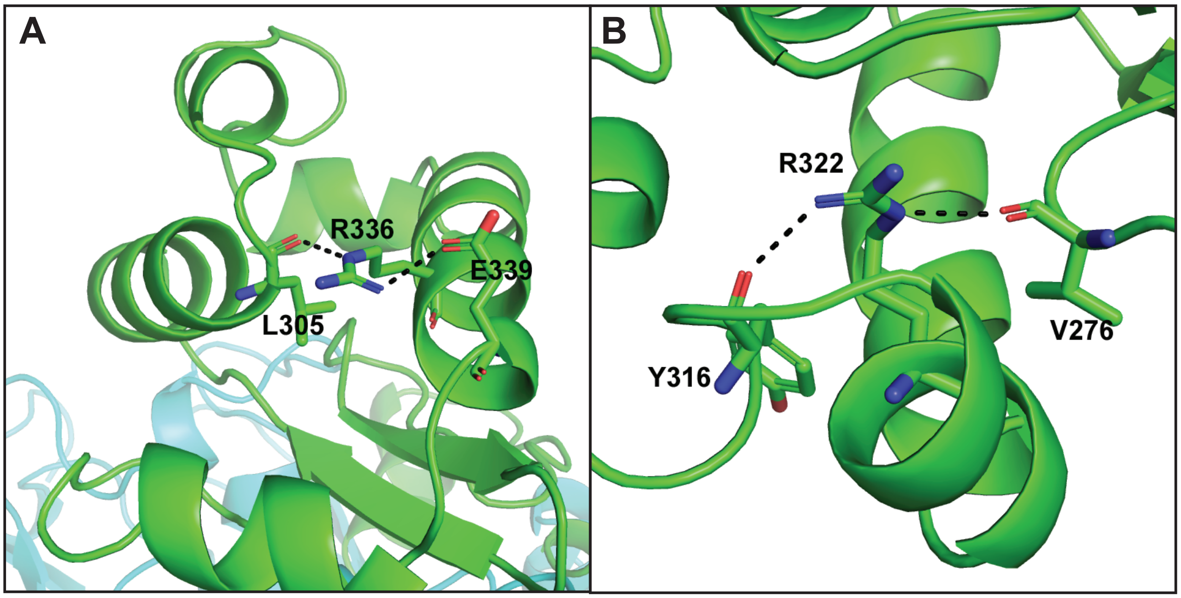

#### Figure S9

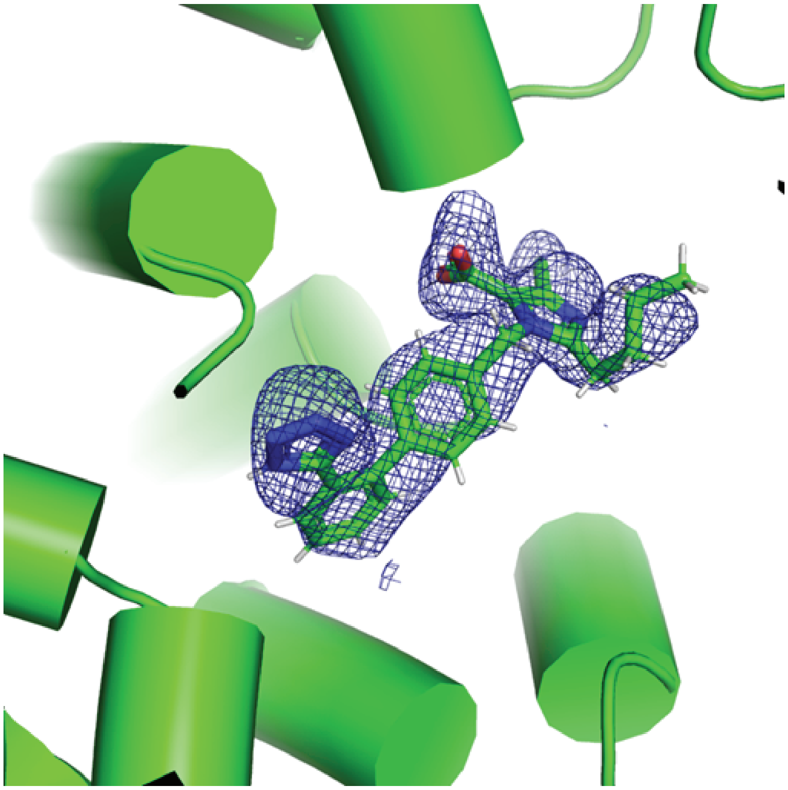

#### Figure S10

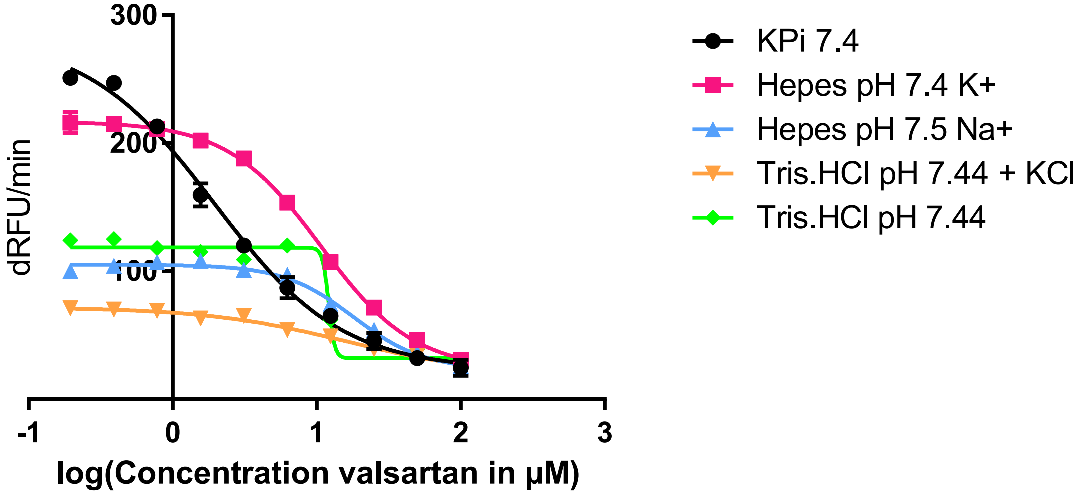

300
200
100
dRFU/min
-1
0
1
2
3
log(Concentration valsartan in µM)
KPi 7.4
Hepes pH 7.4 K+
Hepes pH 7.5 Na+
Tris.HCl pH 7.44 + KCl
Tris.HCl pH 7.44
